# Persister Cells Resuscitate via Ribosome Modification by 23S rRNA Pseudouridine Synthase RluD

**DOI:** 10.1101/678425

**Authors:** Sooyeon Song, Thomas K. Wood

## Abstract

Upon a wide range of stress conditions (e.g., nutrient, antibiotic, oxidative), a subpopulation of bacterial cells known as persisters survive by halting metabolism. These cells resuscitate rapidly to reconstitute infections once the stress is removed and nutrients are provided. However, how these dormant cells resuscitate is not understood well but involves reactivating ribosomes. By screening 10,000 compounds directly for stimulating *Escherichia coli* persister cell resuscitation, we identified that 2-{[2-(4-bromophenyl)-2-oxoethyl]thio}-3-ethyl-5,6,7,8-tetrahydro[1]benzothieno[2,3-d]pyrimidin-4(3H)-one (BPOET) stimulates resuscitation. Critically, by screening 4,267 *E. coli* proteins, we determined that BPOET activates hibernating ribosomes via 23S rRNA pseudouridine synthase RluD, which increases ribosome activity. Corroborating the increased waking with RluD, production of RluD increased the number of active ribosomes in persister cells. Also, inactivating the small RNA RybB which represses *rluD* led to faster persister resuscitation. Hence, persister cells resuscitate via activation of RluD.

## INTRODUCTION

Upon myriad stresses such as antibiotic stress, a sub-population of bacterial cells becomes dormant and multi-stress tolerant (1, 2); these cells are known as persisters. The persister phenotype is not due to genetic change, since upon re-growth, persisters cells behave the same as the original culture. Persistence is relevant in the environment since almost all cells face starvation (3) and relevant in medicine since recurring infections may be the result of regrowth of persister cells (4). The persister sub-population should be distinguished from slow-growing cells such as those in the stationary-phase or those generated by nutrient shifts (5); these slow-growing cells may be distinguished from persisters since the whole population of slow-growing cells are tolerant to antimicrobials whereas the non-growing persister population is a small sub-population (less than 1%) (6). This distinction is critical since tolerant cells utilize alternate sigma factors like RpoS in *Escherichia coli* to redirect gene expression as an active response against stress (7), whereas persisters cease responding and become dormant (5, 8).

To treat persister cell infections, it is important to understand how they form and how they resuscitate. The prevailing view for their formation (6) is that to reduce metabolism, cells activate toxins of toxin/antitoxin (TA) systems (9). The best genetic evidence for this is that deletion of toxins MqsR (10, 11), TisB (12), and YafQ (13) decreases persistence. Moreover, production of non-TAs toxins also increases persistence (14). However, since nutrient deprivation also results in persistence (15), the sub-population of cells may become dormant simply by running out of food. In addition, we have proposed a model whereby the alarmone ppGpp (synthesized as a result of myriad stress conditions), directly creates persister cells via ribosome dimerization, without the need of TA systems (16). Regardless of the mechanism, persistence appears to be an elegantly-regulated response to an unfavorable environment (17).

In regard to resuscitating persister cells, little has been determined about the mechanism. It has been suggested that persister cells resuscitate by inactivating toxins such as TacT acetyltransferase via peptidyl-tRNA hydrolase Pth (18), but this has not been demonstrated. It is established that persister cells revive in response primarily to environmental signals, such as fresh nutrients (rather than stochastically) (19). In addition, persisters revive in an heterogeneous manner, by activating ribosomes; cells increase their ribosome content until a threshold is reached, then they begin to elongate or divide (19). For resuscitation, the persisters sense nutrients by chemotaxis and phosphotransferase membrane proteins, reduce cAMP levels to rescue stalled ribosomes, unhybridize 100S ribosomes via HflX, and undergo chemotaxis toward fresh nutrients (20).

In the present study, to discern additional insights into how ribosomes are active as persister cells resuscitate, we converted the complete *E. coli* population into persister cells so that we could screen for the first time compounds that enhance persister resuscitation. From a 10,000 compound library, we identified that 2-{[2-(4-bromophenyl)-2-oxoethyl]thio}-3-ethyl-5,6,7,8-tetrahydro[1]benzothieno[2,3-d]pyrimidin-4(3H)-one (BPOET) stimulates persister cell waking. Critically, we determined that the mechanism by which BPOET resuscitates persisters is via activation of the 23S rRNA pseudouridine synthase RluD, which is important for ribosome activity. Hence, BPOET stimulates persister resuscitation by activating ribosomes via RluD.

## MATERIALS AND METHODS

### Bacteria and growth conditions

*E. coli* K-12 and its isogenic mutants (**Table 1**) were grown routinely in lysogeny broth (21) (22) at 37°C. BPOET was obtained from ChemBridge (San Diego, CA).

**Table 1.** *E. coli* bacterial strains and plasmids used in this study. Km^R^ and Cm^R^ indicate kanamycin and chloramphenicol resistance, respectively.

| Strains and Plasmids | Features | Source |
| --- | --- | --- |
| BW25113 | Wild type | (39) |
| BW25113 $\Delta rluD$ | $\Delta rluD$ Km<sup>R</sup> | (39) |
| MG1655 | Wild type | (40) |
| MG1655 $\Delta rybB$ | $\Delta rybB$ Km<sup>R</sup> | (41) |
| MG1655 □ ASV | *rrn*bP1::GFP[ASV] | (26) |
| **Plasmids** |  |  |
| pCA24N | Cm<sup>R</sup>; *lacI*<sup>q</sup> | (25) |
| pCA24N_ *rluD* | Cm<sup>R</sup>; *lacI*<sup>q</sup>, P<sub>T5-lac</sub>::*rluD*<sup>+</sup> | (25) |

**Table 2.**
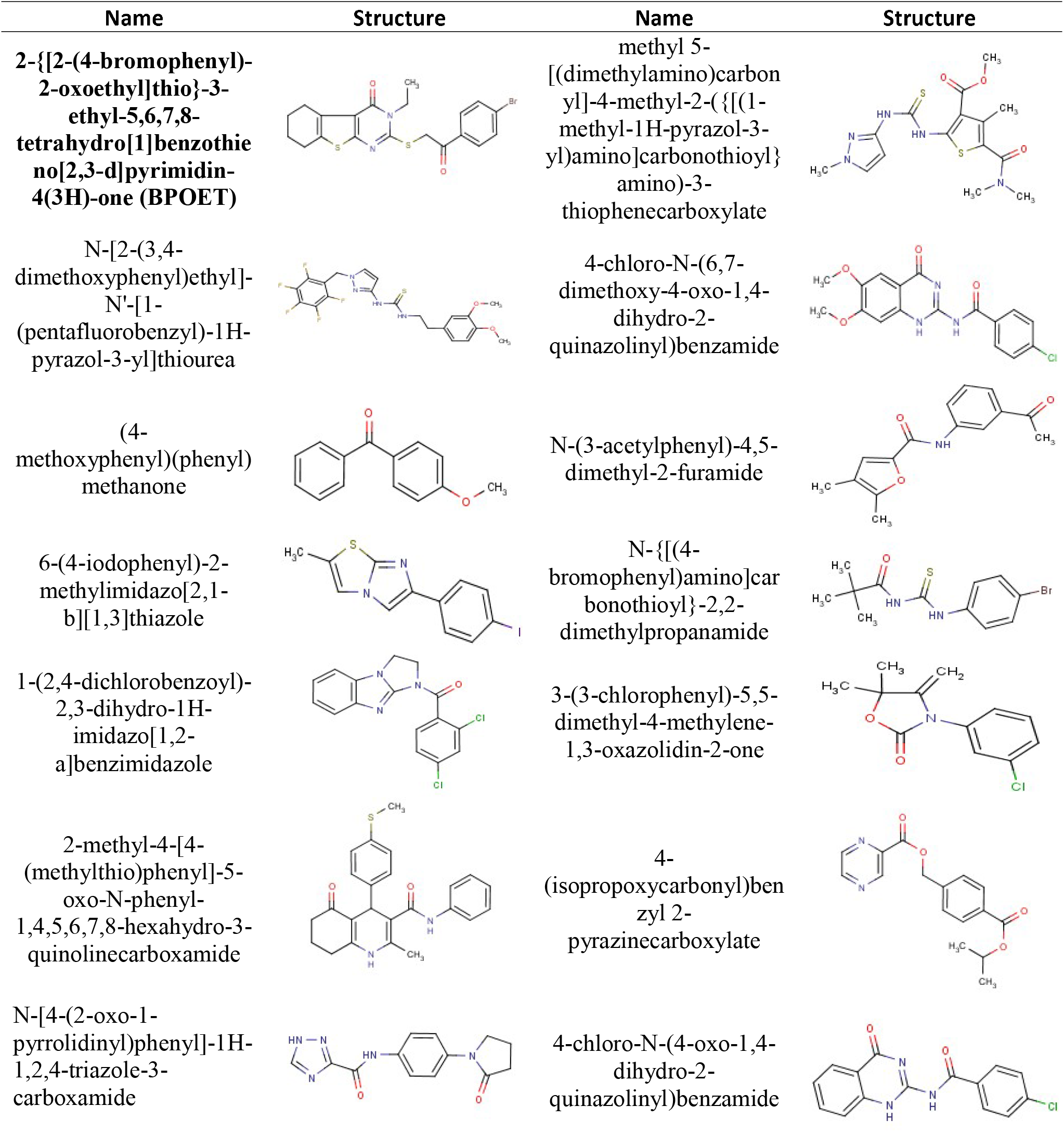
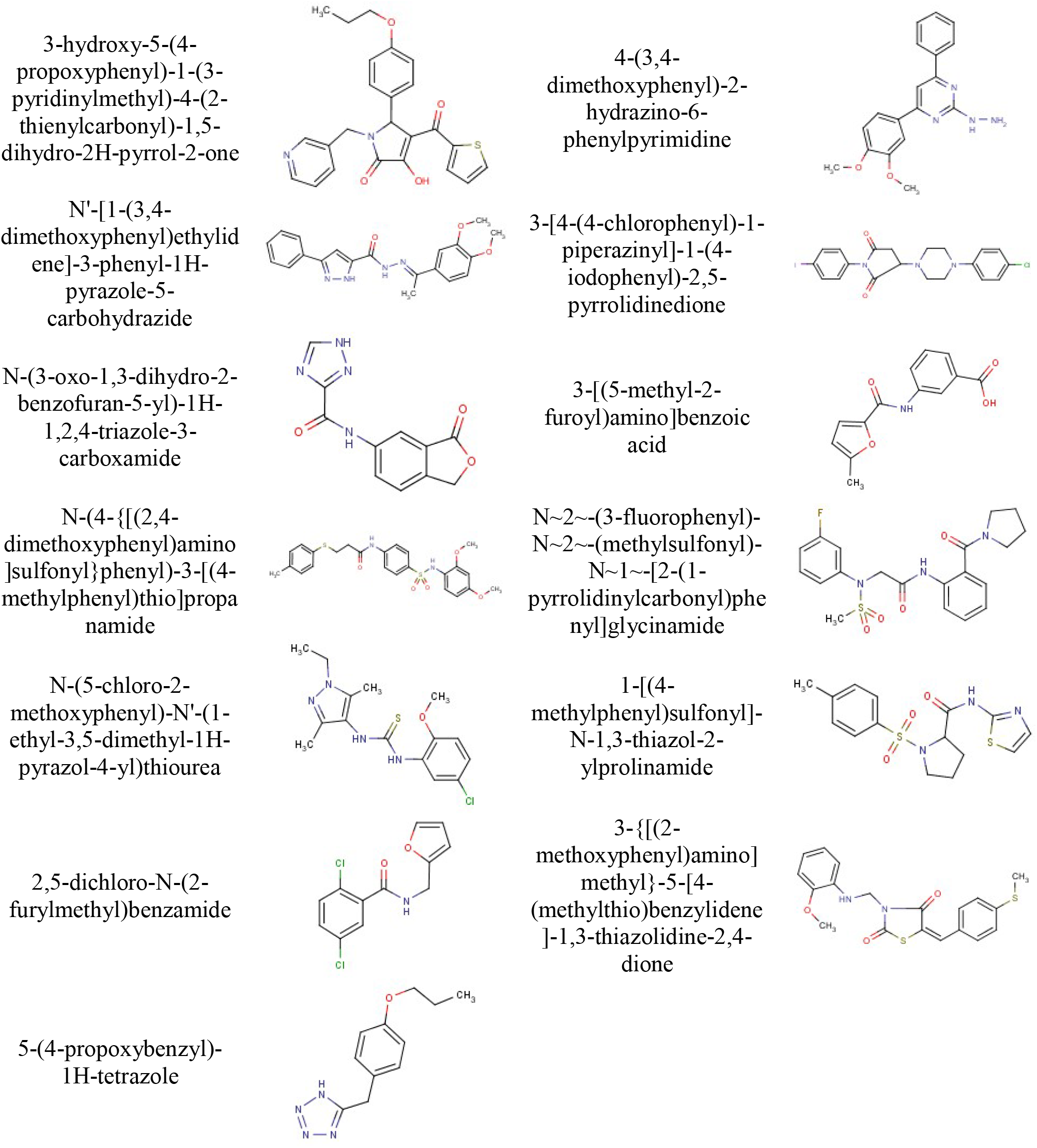
Compounds that resuscitate persister cells and their structures that were identified in the initial screen. Chemical structures are from ChemBridge. BPOET is indicated in bold.

### Persister cells

*E. coli* persister cells were generated (19, 23) by treating exponentially-growing cells (turbidity of 0.8 at 600 nm) with rifampicin (100 μg/mL) for 30 min to stop transcription, centrifuging, and adding LB with ampicillin (100 μg/mL) for 3 h to lyse any non-persister cells. Cells pellets were washed twice with 0.85% NaCl then re-suspended in 0.85% NaCl.

### ChemBridge screen to identify resuscitation compounds

To identify compounds that resuscitate *E. coli* persister cells, the DIVERset Library from ChemBridge (San Diego, CA) containing 10,000 druglike compounds with high pharmacophore diversity was evaluated by adding 4 μL of each compound (final concentration 100 μM, dissolved in DMSO) to 186 μL of LB in 96 well plates and then adding 10 μL of persister cells. The negative control was 2 vol% DMSO. Resuscitation was calculated as the change in turbidity at 600 nm. The compounds that were identified initially were re-tested in M9 minimal medium with 5X alanine (24).

### Pooled ASKA screen to identify resuscitation proteins

To identify proteins responsible for resuscitation, all 4,267 ASKA clones (GFP-) (25) were combined, grown to a turbidity of 2 at 600 nm in LB medium, and their plasmids isolated using a plasmid DNA Mini Kit I (OMEGA Bio-Tek, Inc., Norcross, GA, USA). The pooled ASKA plasmids (1 μL containing 30 ng DNA) were electroporated into 50 μL of *E. coli* BW25113 competent cells, 1 mL LB medium was added, and the cells were grown to a turbidity of 0.5 at 600 nm. Chloramphenicol was added (30 μg/mL) to the culture to maintain the plasmids, and the cells were incubated at 250 rpm to a turbidity of 0.8. Rifampicin followed by ampicillin was added to make persister cells, then the persister cells were washed twice with 1x PBS buffer, contacted with 100 μM BPOET for 2 h in M9 medium that lacked a carbons source, and plated on LB (Cm) agar plates. Faster colony appearance indicated faster persister resuscitation. Plasmids were isolated from the colonies and sequenced using primer pCA24N_F: GCCCTTTCGTCTTCACCTCG.

### Single-cell persister resuscitation

As described previously (19), 5 μL of cell populations consisting of 100% persister cells were added to 1.5% agarose gel pads containing either M9 medium with glucose (0.4 wt%) or alanine (5X) as a carbon source (24), and resuscitation was monitored at 37°C via a light microscope (Zeiss Axio Scope.A1, bl_ph channel at 1000 ms exposure).

### Active 70S ribosome assay

The GFP signal of resuscitating persisters of *E. coli* K-12 MG1655-ASVGFP (26) with RluD was monitored using a fluorescence microscope (Zeiss Axioscope.A1, bl_ph channel at 1,000 ms exposure and GFP channel at 10,000 ms exposure). *E. coli* K-12 MG1655-ASVGFP produces an unstable variant of GFP (half-life less than 1 h) under the control of the 16S rRNA ribosomal promoter *rrnb*P1 (26).

## RESULTS & DISCUSSION

### BPOET resuscitates *E. coli* persister cells

To identify compounds that resuscitate *E. coli* persister cells, we first increased by 10^5^-fold the persister cell population by pre-treating with rifampicin to cease metabolism by stopping transcription followed by ampicillin treatment to kill any remaining non-persister cells (23). In this way, nearly 100% of bacterial cell population was converted into persister cells. Hence, we were able to both screen for compounds that more rapidly resuscitate persister cells as well as confirm our hypotheses via single-cell microscopy. The persister cells generated in this way have been (i) confirmed eight ways (19), (ii) used to determine that persister cells wake via ribosome activation (19) and chemotaxis (20), (iii) used to show that the cells capable of resuscitation in a viable but not culturable population are equivalent to persister cells (15), (iv) used to identify compounds that kill persister cells (27), and (v) used to show that the alarmone ppGpp directly creates persister cells by stimulating ribosome dimerization (16). In addition, our method to generate a high population of persister cells has been utilized by at least six independent groups (28–33).

Using 96-well plates, the persister cells (10 μL) were added to 190 μL of LB containing one each of the 10,000 compounds of the DiverSet library dissolved in dimethyl sulfoxide (100 μM final concentration), and growth was monitored via the change in turbidity for up to 48 h. Starting at a turbidity of 0.05, a 140-fold increase in growth was possible (maximum final turbidity of 0.69). **Table 1** shows the 27 compounds that were identified that stimulated persister cell resuscitation relative to the negative control of dimethyl sulfoxide. Upon confirming the results of these initial hits in minimal alanine medium, we found BPOET (100 μM) was most effective and increased persister cell waking by 44-fold in 96-well plates based on the increases in turbidity as well as found that BPOET increases the waking of single persister cells by 4-fold (**Figure 1**, **Table S1**). Hence, we focused on this compound.

**Fig. 1.**
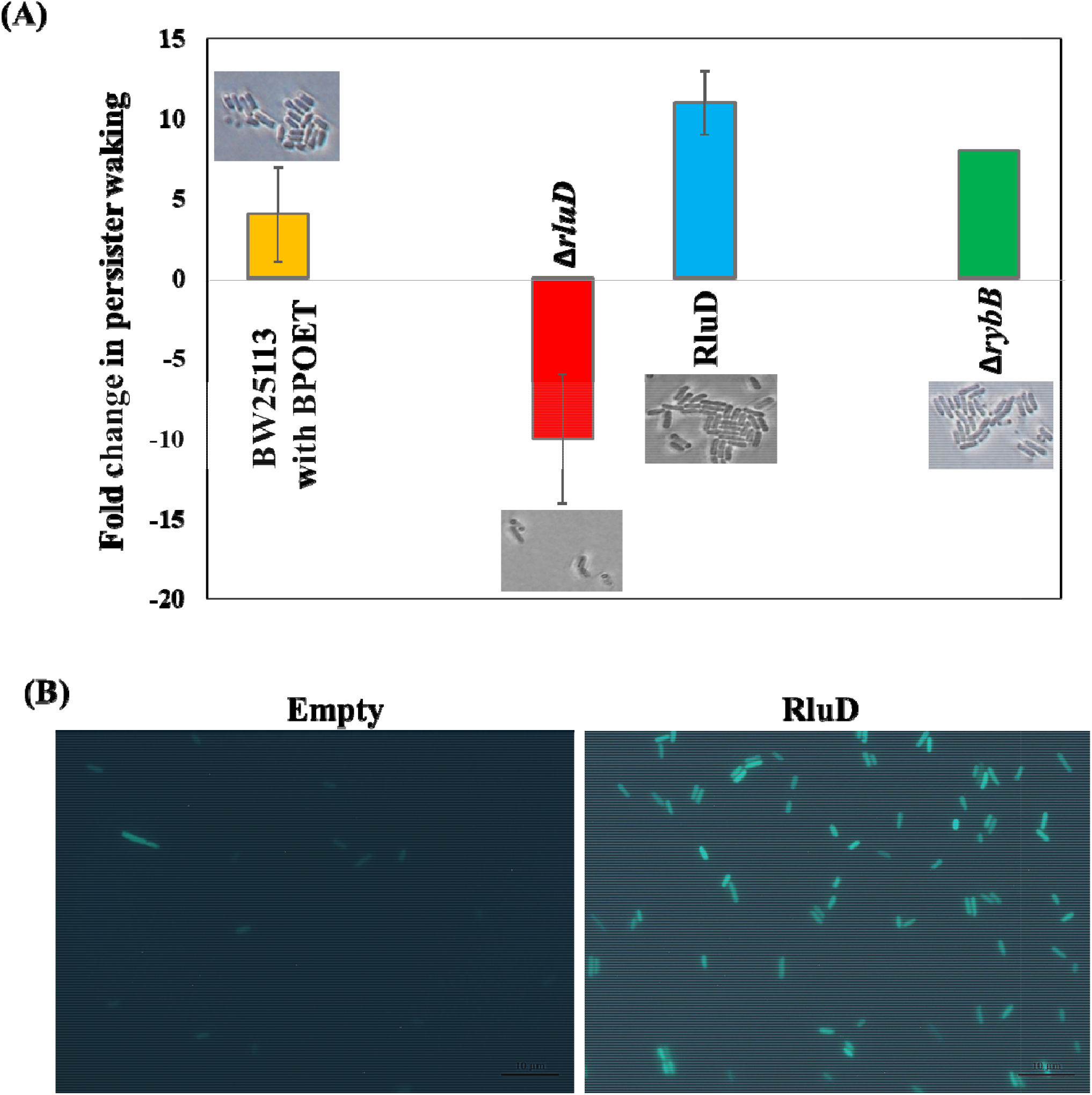
RluD increases persister resuscitation by increasing ribosomes for resuscitation. (**A**) Single-cell persister resuscitation as determined using light microscopy (Zeiss Axio Scope.A1). The total and waking number of persister cells are shown in **Table S1**. Microscope images for waking cells are shown in **Fig. S1**. The fold-change in resuscitation is relative to BW25113 with DMSO for BW25113 with BPOET, relative to BW25113 for the Δ*rluD* mutant, relative to BW25113/pCA24N for the strain producing RluD from pCA24N plasmid in BW25113, and relative to MG1655 for Δ*rybB*. M9 glucose (0.4%) agarose gel pads were used for all the strains except BW25113 with BPOET where M9 alanine (5X) agarose gel pads including 100 μM of BPOET or DMSO were used. The results are the combined observations from two independent experiments after 6 h for the BW25113 with BPOET, after 4 h for BW25113 and its deletion mutants, and after 6 h for cells harboring pCA24N and its derivatives as well as for MG1655 and MG1655 Δ*rybB*. Error bars indicate standard deviations. (**B**) Active 70S ribosomes in single persister cells for MG1655-ASV/pCA24N-rluD (“RluD”) vs. MG1655-ASV/pCA24N (“Empty”). Cells are shown on agarose gel pads at time 0 for resuscitation; i.e., after the formation of persister cells. Representative results from three independent cultures are shown.

### BPOET resuscitates *E. coli* persister cells by modifying ribosomes

To determine how BPOET resuscitates persister cells, we pooled the 4,267 ASKA clones in which each *E. coli* protein is produced from plasmid pCA24N, produced persister cells carrying these plasmids, contacted with 100 μM BPOET, plated the cells, and chose the largest colonies that formed on LB plates. Our rationale was that any pathway stimulated by BPOET would be even more active if the number of rate-limiting proteins in that pathway were increased, and cells that wake first would form colonies faster.

Using this approach, we identified five proteins whose production increased resuscitation: RluD, YjiK, SrlR, Smf, and YeeZ. These proteins are related to contacting with BPOET since addition of the diluent DMSO alone and sequencing larger colonies did not identify these five proteins but instead identified TmcA, a tRNA^Met^ cytidine acetyltransferase, which is a general factor required for translation that likely led to larger colony sizes with the diluent. Of the proteins related to BPOET, only RluD (23S rRNA pseudouridine synthase) and SrlR (represses the *gut* operon for glucitol metabolism) have been characterized; we focused on RluD because it is related to ribosomes, and we have shown inactivating ribosomes causes persistence (23) and activating ribosomes resuscitates persister cells (16, 19, 20). RluD is involved in the synthesis and assembly of 70S ribosomes as well as their function based on its post-transcriptional modification of 23S rRNA to form three pseudouridine (5-ribosyl-uracil) nucelosides at positions 1911, 1915, and 1917 (34). In pseudouridine, uracil is attached via a carbon-carbon bond to the sugar base rather than through a carbon-nitrogen bond. The 23S rRNA pseudouridines increase the stability of the tertiary structure of 23S rRNA and are located in a stem loop structure that is involved in peptidyltransferase and interacts with mRNA, tRNA, 16S rRNA, and ribosome release factor. Hence, RluD is responsible for efficient ribosome function (34).

### RluD enhances persister cell resuscitation

To explore further the role of RluD and persister resuscitation, we utilized single cell studies since persister cells are heterogeneous (19) and wake with different frequencies (which would be missed if we monitored planktonic populations). We found that deleting *rulD* reduces the frequency of single-cell persister resuscitation dramatically (11-fold) compared to the isogenic wild-type strain on minimal glucose agarose gel pads (**Fig. 1A**, **Table S1**, **Fig. S1**). In addition, no colonies were found on M9 glucose agar plates after inactivating RluD (**Fig S2**), confirming that persister cells are severely challenged in resuscitation without RluD. Corroborating these two results with the *rluD* mutant, production of RluD increased the frequency of waking by 11-fold on glucose medium (**Fig. 1A**, **Table S1**, **Fig S1**) and increased waking on rich medium (results not shown). In addition, the *rluD* deletion has no effect on persister cell formation (**Fig S2**). Therefore, RluD stimulates persister cell resuscitation but does not affect persister formation.

### RluD increases active ribosomes for resuscitation

Using a GFP reporter that indicates the number of 70S ribosomes in individual persister cells (19), we found that producing RluD before making persister cells makes 85 ± 6% of the cells have high ribosome fractions compared to not producing RluD (**Fig. 1B**). The GFP reporter indicates transcription of *rrsB* (16S rRNA), *gltT* (tRNA-glu), *rrlB* (23S rRNA) and *rrfB* (5S rRNA); hence, it indicates production of the three major rRNA building blocks. Although this is not a direct observation of 70S ribosomes, this method is a suitable proxy for the number of ribosomes based on measurement of rRNA concentrations and has been used frequently (19, 35–37), and we have verified its use by isolating ribosomes and comparing GFP fluorescence (19). Hence, the increased persister cell resuscitation with RluD is directly due to the increase in active (70S) ribosomes of persister cells.

### RybB antagonizes persister cell resuscitation

Since the small RNA RybB represses RluD (38), we investigated its impact on persister resuscitation. As expected, we found that deletion of *rybB* increases the frequency of persister cell waking by 8-fold (**Fig. 1A**, **Table S1**, **Fig. S1**).

In summary, the results presented here demonstrate that ribosomes may be activated for specific cell cycles such as recovery from dormancy. Specifically, by screening for compounds for the first time that enhance persister cell resuscitation, we have (i) determined that ribosomes are modified by RluD as cells resuscitate and resume ribosome activity, (ii) identified a novel compound, BPOET, that activates persister cells, and (iii) linked small RNAs to persistence. Hence, these results extend our understanding of how persister cells are activated which has a far-reaching impact in that all bacteria cope with nutrient stress and become dormant.

## ACKNOWLEDGEMENTS

This work was supported by funds derived from the Biotechnology Endowed Professorship at the Pennsylvania State University. We are grateful for the *rybB* mutant provided by Dr. Gisela Storz. The authors have no conflicts of interest.

## SUPPORTING INFORMATION

**Table S1.**
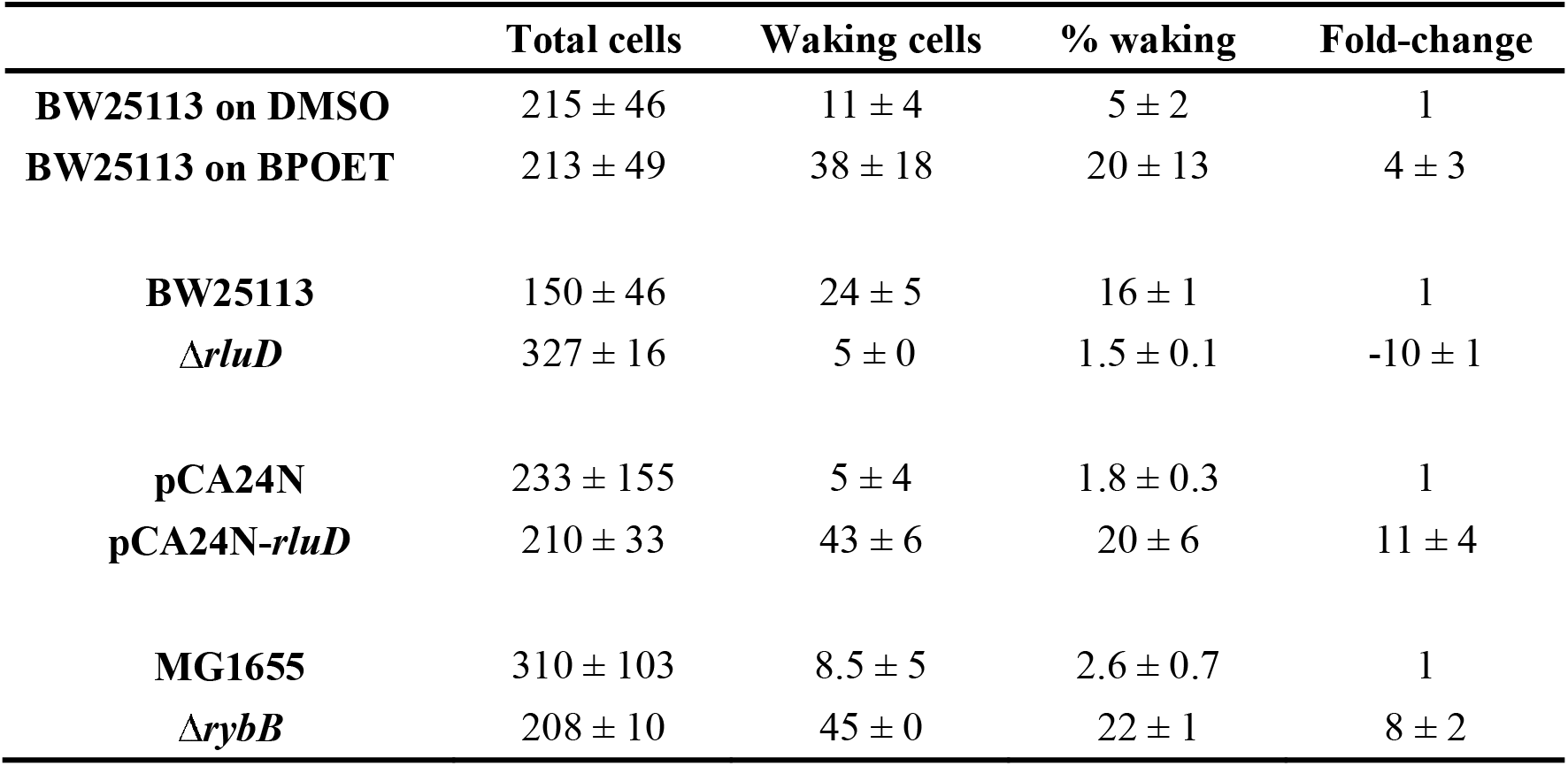
Single persister cell resuscitation. Single-cell persister resuscitation as determined using light microscopy (Zeiss Axio Scope.A1) using agarose gel pads. Microscope images are shown in **Fig. S1**. The fold-change in resuscitation is relative to BW25113 with DMSO for BW25113 with BPOET, relative to BW25113 for the Δ*rluD*, relative to BW25113/pCA24N for the strain producing RluD from pCA24N in BW25113, and relative to MG1655 for Δ*rybB*. M9 glucose (0.4%) agarose gel pads were used for all the strains except BW25113 with BPOET where M9 alanine (5X) agarose gel pads including 100 μM of BPOET or DMSO were used. The results are the combined observations from two independent experiments after 6 h for the BW25113 with BPOET and DMSO, after 4 h for BW25113 and its deletion mutants, and after 6 h for cells harboring pCA24N and its derivatives as well as for MG1655, and Δ*rybB*. Standard deviations are shown, and each strain was visualized at 14 positions.

**Table S2.** Active 70S ribosomes in single persister cells for MG1655◻ASV/pCA24N-rluD (“pCA24N-*rluD*”) vs. MG1655◻ASV/pCA24N (“pCA24N”). Single-cell persister resuscitation as determined using light microscopy (Zeiss Axio Scope.A1) using agarose gel pads with 0.4% glucose. Microscope images are shown in **Fig. 1B**. The fold-change in resuscitation is relative to MG1655◻ASV/pCA24N for MG1655◻ASV/pCA24N-rluD.

|  | pCA24N-rluD | pCA24N |
| --- | --- | --- |
| **Total cells** | 140 ± 66 | 112 ± 4 |
| **High intensity cells** | 120 ± 64 | 25 ± 3 |
| **Waking %** | 85 ± 6 | 22 ± 2 |
| **Fold-change** | 3.8 ± 0.4 | 1 |

**Figure S1.**
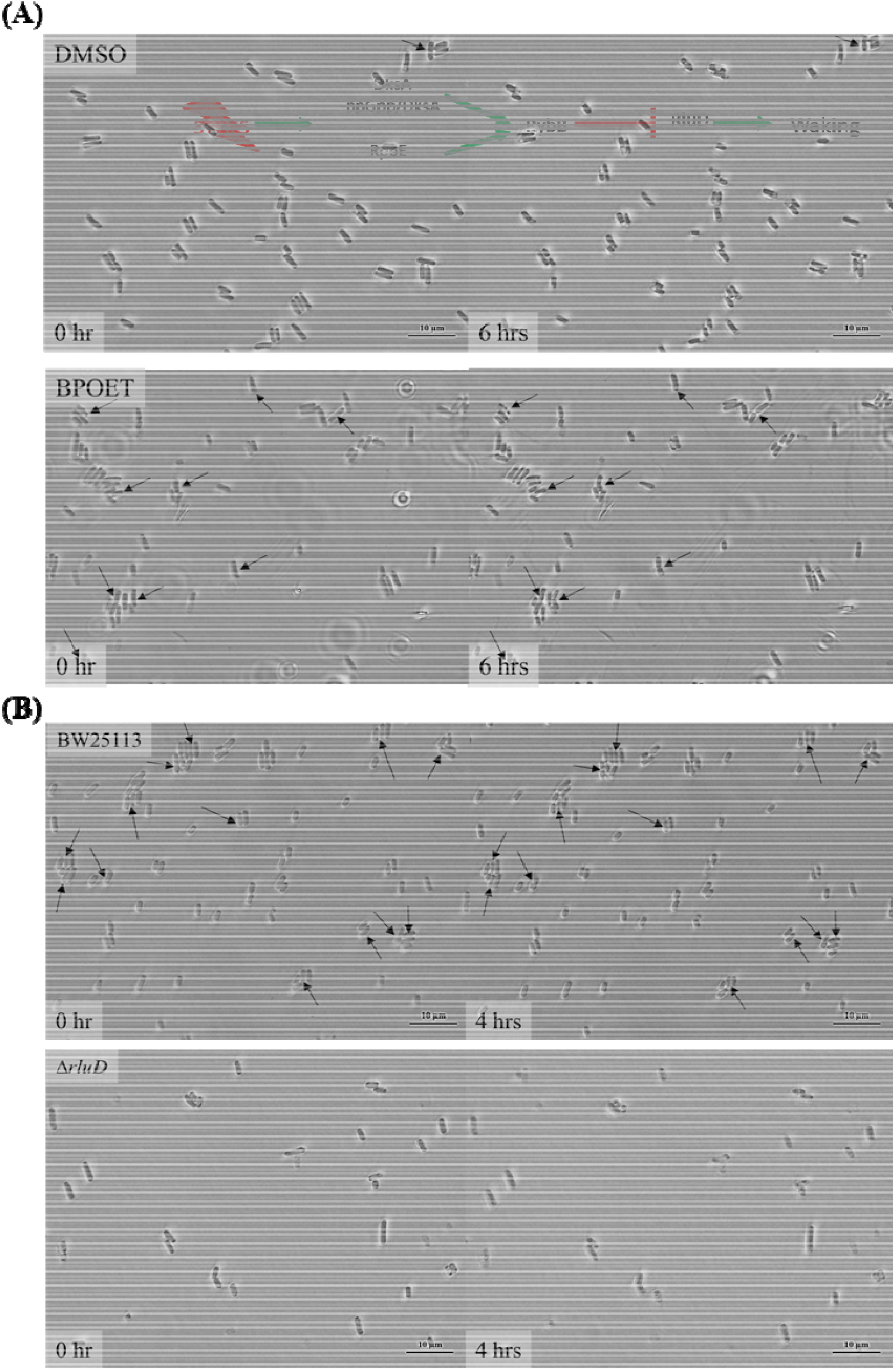
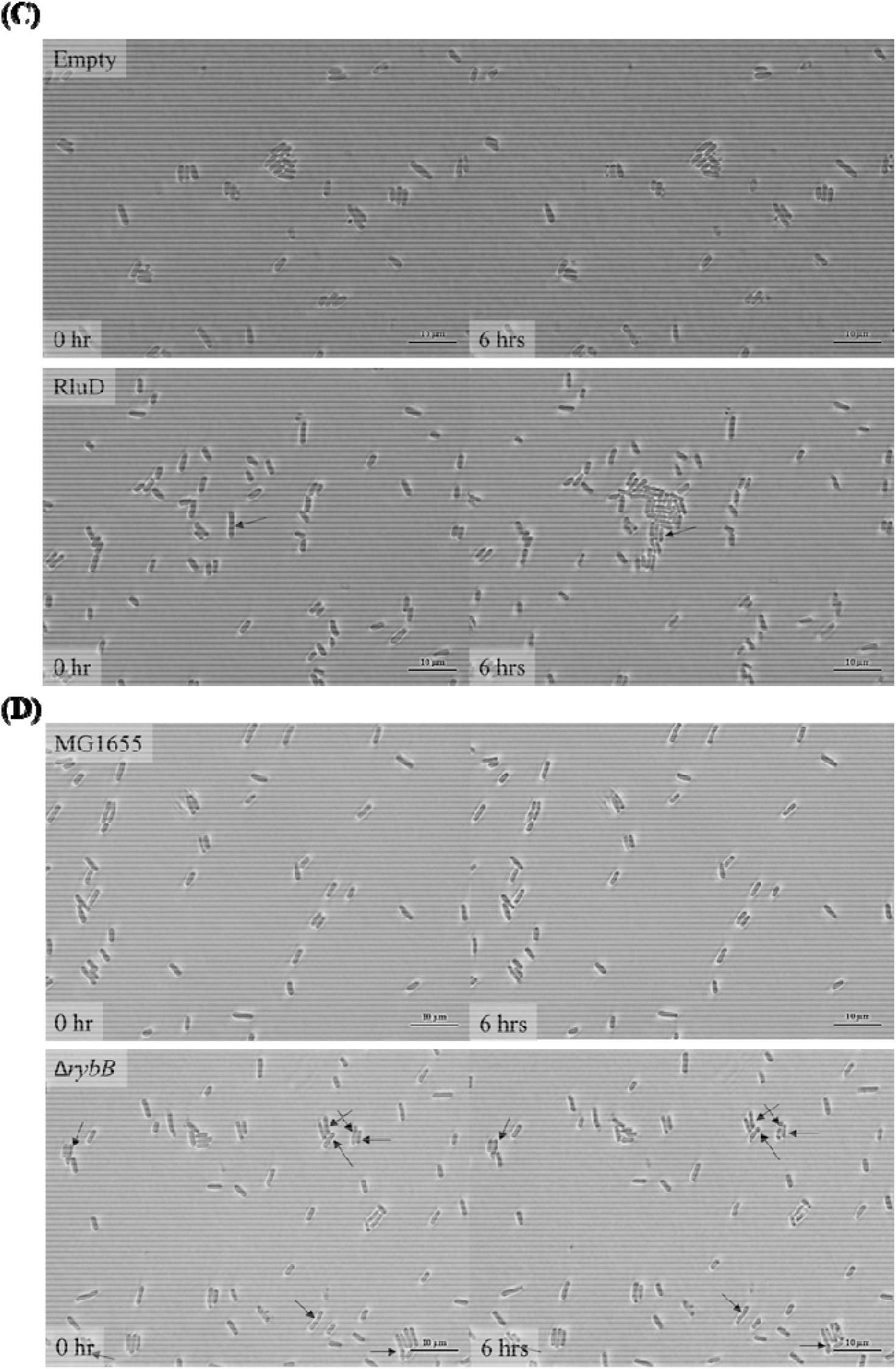
Single persister cell waking. Persister cells of (**A**) BW25113 with DMSO (upper panel), and BPOET (lower panel) on M9 5X Ala agarose gel pads containing DMSO and BPOET (100 μM) after 6 h, (**B**) BW25113 (upper panel) and BW25113 Δ*rluD* (lower panel) after 4 hours on M9 0.4% glucose agarose gel pads, (**C**) BW25113/pCA24N (“Empty”), and BW25113/pCA24N-*rluD* (“RluD”) after 6 h on M9 0.4% glucose agarose gel pads, and (**D**) MG1655, and MG1655 Δ*rybB* after 6 h on M9 0.4% glucos agarose gel pads. Black arrows indicate cells that resuscitate. Scale bar indicates 10 μm. Representative results from two independent cultures are shown.

**Figure S2.**
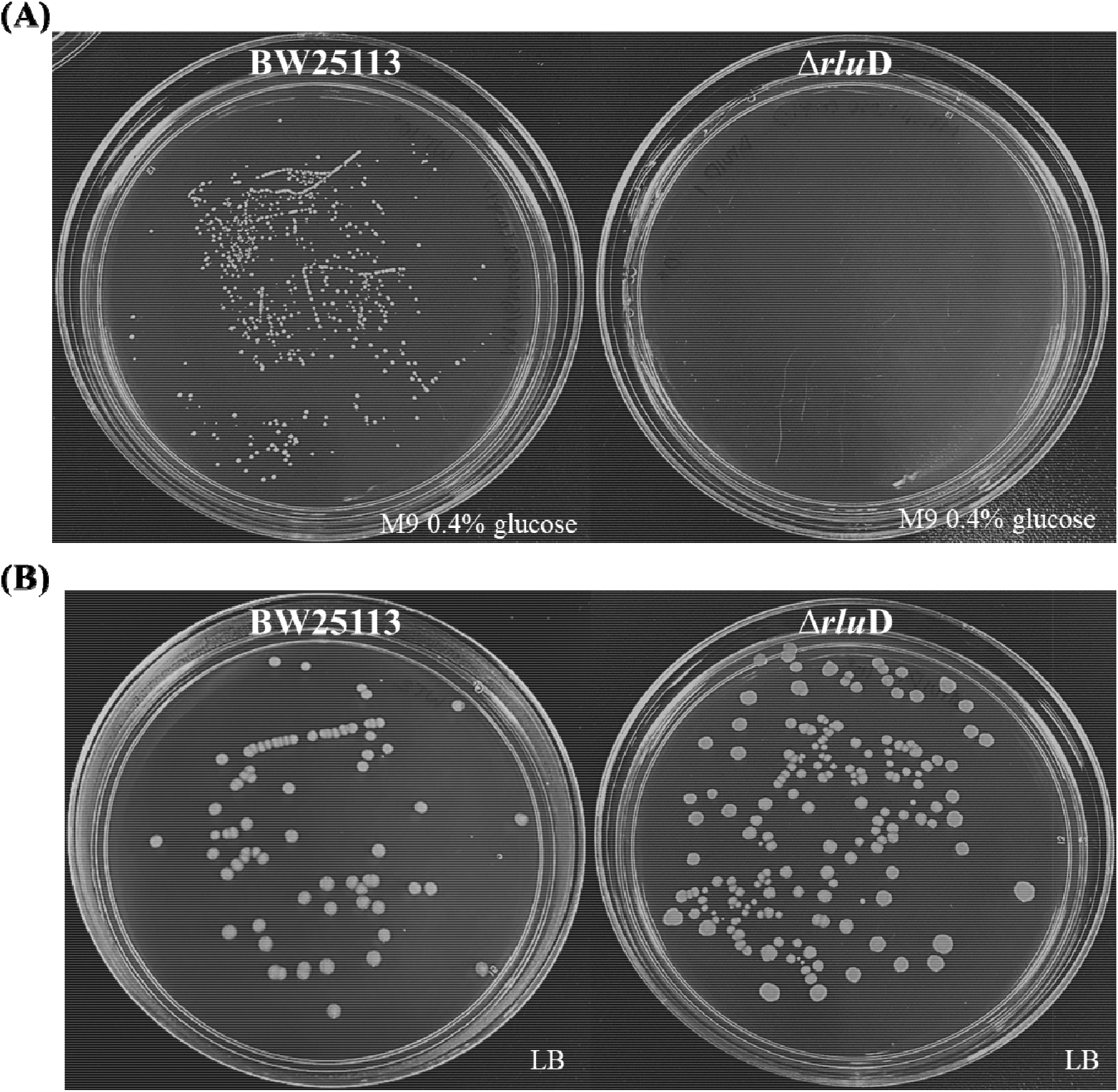
Inactivating RluD eliminates persister cell waking on minimal glucose medium but doe not affect the number of persister cells that are formed. (**A**) Resuscitation of wild type BW25113 and BW25113 Δ*rluD* persister cells at 37 °C on M9 0.4% glucose agar plates for three days. (**B**) Colonies formed in one day at 37 °C on LB agar plates indicating the number of persister cells for BW25113 and the isogenic Δ*rluD* mutant. One representative plate of two independent cultures is shown.

